# Pitfalls and opportunities for applying PEER factors in single-cell eQTL analyses

**DOI:** 10.1101/2022.08.02.502566

**Authors:** Angli Xue, Seyhan Yazar, Drew Neavin, Joseph E. Powell

**Affiliations:** Garvan-Weizmann Centre for Cellular Genomics, Garvan Institute of Medical Research, Sydney, NSW, 2010, Australia; School of Medical Sciences, University of New South Wales, Sydney, NSW, 2052, Australia; UNSW Cellular Genomics Futures Institute, University of New South Wales, Sydney, NSW, 2052, Australia

**Keywords:** Single-cell RNA-seq, Pseudo-bulk, Latent variable, PEER factors, Normalisation, eQTL mapping

## Abstract

Using latent variables in gene expression data can help correct spurious correlations due to unobserved confounders and increase statistical power for expression Quantitative Trait Loci (eQTL) detection. Probabilistic Estimation of Expression Residuals (PEER) is a widely used statistical method that has been developed to remove unwanted variation and improve eQTL discovery power in bulk RNA-seq analysis. However, its performance has not been largely evaluated in single-cell eQTL data analysis, where it is becoming a commonly used technique. Potential challenges arise due to the structure of single-cell data, including sparsity, skewness, and mean-variance relationship. Here, we show by a series of analyses that this method requires additional quality control and data transformation steps on the pseudo-bulk matrix to obtain valid PEER factors. By using a population-scale single-cell cohort (OneK1K, *N* = 982), we found that generating PEER factors without further QC or transformation on the pseudo-bulk matrix could result in inferred factors that are highly correlated (Pearson’s correlation *r* = 0.626∼0.997). Similar spurious correlations were also found in PEER factors inferred from an independent dataset (induced pluripotent stem cells, *N* = 31). Optimization of the strategy for generating PEER factors and incorporating the improved PEER factors in the eQTL association model can identify 9.0∼23.1% more eQTLs or 1.7%∼13.3% more eGenes. Sensitivity analysis showed that the pattern of change between the number of eGenes detected and PEER factors fitted varied significantly for different cell types. In addition, using highly variable genes (e.g., top 2000) to generate PEER factors could achieve similar eGenes discovery power as using all genes but save considerable computational resources (∼6.2-fold faster). We provide diagnostic guidelines to improve the robustness and avoid potential pitfalls when generating PEER factors for single-cell eQTL association analyses.

## Background

Inferring latent variables that explain the variation in the gene expression data has been an essential step for expression Quantitative Trait Loci (eQTL) analyses. It can be used to identify the unobserved confounding effects and potential cellular phenotypes (e.g., transcription factor or pathway activation). Standard methods of inferring latent variables include principal component analysis (PCA)^1^, surrogate variable analysis (SVA)^2^, and Probabilistic Estimation of Expression Residuals (PEER)^3,4^. PEER is a method that implements a Bayesian framework to estimate the latent variables and jointly learn the contribution to the gene expression variability from genotype, known factors, and hidden factors. The inferred factors (i.e., PEER factors) can be applied to increase the power of eQTL discovery. This method was introduced in 2010 and is widely used in bulk eQTL analyses^5-8^, and recently the emerging field of single-cell pseudo-bulk eQTL analysis^9-12^.

As the scale of single-cell RNA-sequencing (scRNA-seq) studies has rapidly grown, eQTL analyses that use pseudo-bulk analysis approaches in scRNA-seq have started to emerge. Pseudo-bulk refers to the aggregation of the gene expression profiling of all cells from one sample into a single pseudo-sample; thus, the data structure will be assimilated into the bulk RNA-sequence data. However, due to the nature of scRNA-seq data structures, the bulk expression matrix and single-cell expression matrix can be very different. There are three main differences between single-cell and bulk RNA data: matrix sparsity, distribution normality or skewness, and mean-variance dependency. First, since the scRNA-seq matrix is sparse and most elements are zero, the pseudo-bulk gene expression matrix also contains many zero values. Second, the underlying distribution of gene expression across cells follows either negative binomial (NB) or zero-inflated NB (ZINB) distributions^13^. Therefore, most inter-individual distributions of mean gene expression in pseudo-bulk are non-normal and heavily right-skewed. Third, mean-variance dependency exists between the intra-individual mean and variance due to the characteristics of the underlying distribution, and such relationships could be retained in the pseudo-bulk data. These features of pseudo-bulk data may violate the assumptions of the PEER method.

Consequently, the inferred PEER factors could suffer from biases or spurious correlations with each other, which can lead to the problematic interpretation of the factors themselves and compromise the discovery power of pseudo-bulk eQTL association. Moreover, it is unclear how many PEER factors should be fitted in the eQTL association model to maximise the detection power in pseudo-bulk data. Previous bulk eQTL analysis either chose a fixed number^7^ or a certain threshold based on the sample size^5,8^. Some studies have run sensitivity tests^5,6,8^, but such optimisation has not been systematically evaluated for single-cell data at the population-scale level.

Here, we identify some common scenarios where pitfalls occur and how they can be avoided with data-driven approaches. To help with the future application of PEER factors to single-cell RNA-sequence data, we propose guidelines for the quality control and scaling of the pseudo-bulk expression matrix, diagnosing and troubleshooting the inferred hidden determinants, and the way to select the optimal number of PEER factors to improve the eQTL discovery.

## Results and Discussion

Using three independent scRNA-seq datasets, we investigated how PEER factors behave under different quality control scenarios and transformations: one from peripheral blood mononuclear cells (PBMCs, *N* = 982) and the others from fibroblasts and induced pluripotent stem cells (iPSCs)^10^ (*N* = 79 and 31). The PBMC data were from the OneK1K cohort^12^, a population-scale scRNA-seq dataset containing ∼1.2 million immune cells collected from 982 donors (Methods). This dataset was quality controlled (QC), normalised and variance stabilised at the single-cell level by *sctrasnform*^14^, and classified into 14 cell types by *scPred*^15^ (**Methods**). To construct a pseudo-bulk expression matrix for each cell type, the gene expression level per individual was calculated as the intra-individual mean counts across cells. We first generated PEER factors while including sex, age, and six genotype PCs as covariates. Using CD4_NC_ cells as an example (**Figure 1**, other cell types shown in **Supp Figure 1**), we observe strong correlations among PEER factors, many of which were nearly equivalent (**Figure 1A**). For instance, while most known covariates are not correlated (Pearson’s *r* = -0.04 ∼ 0.06, except -0.13 between PC3 and PC4; **Supp Figure 2**), the first and second PEER factors have a modest correlation (Pearson’s *r* = 0.20). However, PEER factors 5-7 have pair-wise correlations equal to 1. Although the hidden factor model of PEER allows for non-orthogonal components, the mean of the pair-wise Pearson’s *r* across the first 10 PEER factors were all larger than 0.5 in all 14 cell types, suggesting that PEER factors are overfitted.

**Figure 1.**
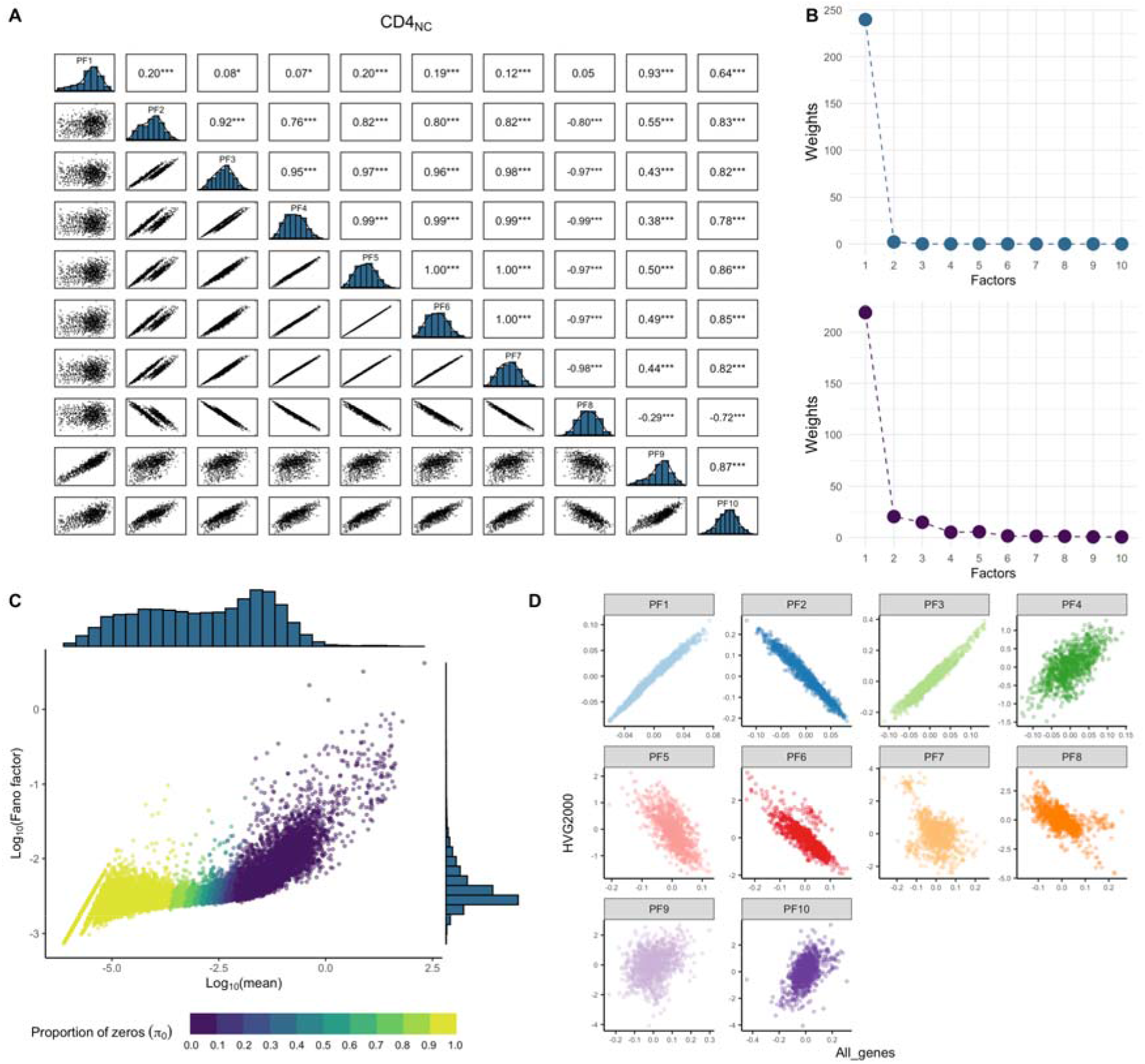
Correlation among inferred PEER factors and global intra-individual mean-variance dependence. **A**, Pair-wise correlation plot among the first 10 PEER factors generated from pseudobulk expression in CD4_NC_ cells without any quality control (option #1). The upper triangle panel shows the pair-wise estimates of Pearson’s correlation, and the bottom triangle panel shows the pair-wise scatter plot between the PEER factors. The diagonal panel shows the distribution of each PEER factor. Significance of correlation test is annotated by * *p*-value ≤ 0.05, ** ≤ 0.01, *** ≤ 0.001. **B**, Diagnostic plot of the factor weights without any further quality control on the pseudo-bulk matrix (option #1, upper panel) and option #11 QC (lower panel). **C**, Relationship between intra-individual pseudo-bulk mean and Fano factor per gene. Both axes are Log10 transformed. The colour of the dots indicates the proportion of zero expression across individuals () for each gene. **D**, scatter plot of first 10 PEER factors generated from all genes against those from top 2000 highly variable genes (option #11 vs option #12).

Additionally, we found that the variance explained by the first PEER factor was overwhelmingly more significant than the rest of the PEER factors, where the latter’s contributions seem negligible (upper panel in **Figure 1B** and **Supp Figure 1B**). This is consistent with the observation that there is a mean-variance dependency in the pseudo-bulk expression level of each gene (**Figure 1C**). Hence, the highly expressed genes inherently contribute much more variation than other genes. Due to the sparsity in scRNA-seq data, there is a certain proportion of genes whose intra-individual expression is zero (**Figure 1C**); therefore, regardless of the transformation or normalisation methods that are used, the intra-individual distribution of these genes will be strongly right-skewed and violate the normality assumption of PEER (see examples in **Supp Figure 3**).

To alleviate the impact of these properties, we tested different options in combinations (13 options in total) to generate PEER factors: (1) excluding the genes with zero expression in more than a certain % of the individuals for all analyses (i.e., π_0_ ≥ 0.9 0r 1); (2) log(x+1) transformation; (3) standardisation, which scales the distribution to mean = 0 and standard deviation = 1; (4) Rank-Inverse Normal Transformation (RINT); (5) Selecting the top 2,000 highly variable genes (HVGs, ranked by the Fano factor, e.g. variance-to-mean ratio) to generate the PEER factors (**Methods**). The results show that the correlation among PEER factors was still high when genes with high π_0_ were excluded, and gene expression was log(x+1) transformed (options #1-5, **Figure 2A**). Among options #6-11, option #7 (standardization + π_0_ ≥ 0.9 excluded) and option #11 (log(x+1) + standardization + π_0_ ≥ 0.9 excluded) had the lowest mean pair-wise correlation between independent PFs (**Figure 2A** and Supp **Figure 4**). We identified option #11 as the optimal performing approach because the skewness of genes was lower than option #7 (median skewness for all genes is 0.86∼3.8 vs 0.90∼5.12 across 14 cell types). We also tried to generate PEER factors using the top 2,000 HVGs (options #12-13 in **Figure 2A-B**). The PEER factors generated from all genes are highly correlated with those from the top 2,000 HVGs (**Figure 1D** and **Supp Figure 1D**), highlighting that the HVGs could explain most of the variation that was explained when using all the genes and reduce the runtime from 46.2 mins to 7.4 mins on average for different cell types (**Figure 2B**).

**Figure 2.**
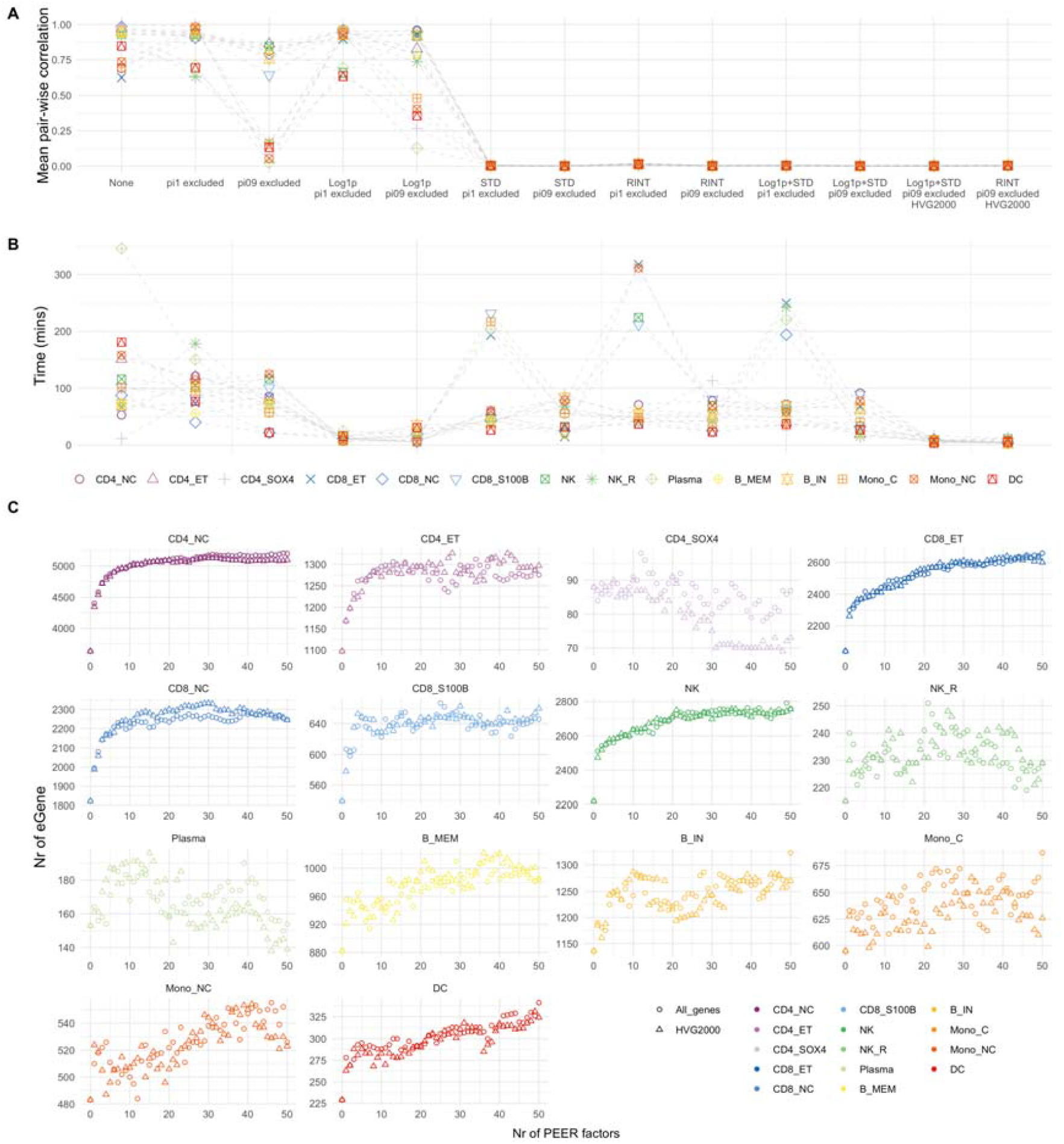
Performance of different QC options on generation of PEER factors and sensitivity test for eGene detection. **A**, The mean pair-wise correlation among the first 10 PEER factors. Each colour and shape represent a specific cell type. **B**, Time to generate 50 PEER factors by different quality control options on the pseudo-bulk matrix. **C**, The x-axis denotes the number of PEER factors fitted as covariates in the association model. The y-axis represents the number of eGenes with at least one eQTL at local FDR < 0.05. The shape of each scatter point indicates whether using all genes or the top 2000 highly variable genes to generate PEER factors (both excluded genes with π_0_ > 0.9, log(x+1) transformed and standardised).

Next, we sought to investigate how PEER factors generated from different strategies affect the power of eQTL discovery. We calculated PEER factors using all genes or the top 2,000 HVGs (both pre-excluded genes with π_0_ ≥ 0.9) and compared the number of eGenes (at least associated with one significant eQTL) identified when incrementally fitting PEER factors as covariates from 0 to 50. Notably, the pattern of change in the number of eGenes varied across different cell types (**Figure 2**). For CD4_NC_ cells, the number of eGenes continually increased until reaching an asymptote of around 30, while CD4_SOX4_ cells reached a peak between 10-15 and decreased as more factors were included. We also show that the pattern of change in eGenes discovery power was consistent regardless of using all genes or the top 2,000 HVGs (**Figure 2C**), and the gains of made in number of detected eGenes were also similar (**Supp Table 1**). These consistencies reaffirmed that using the top 2,000 HVGs captures most of the latent variation that can be explained by all genes in this dataset. We also compared the number of eQTLs/eGenes when PEER factors were generated without QCs or QC option #11 on the pseudo-bulk matrix. The QC option #11 can identify 9.0∼23.1% more eQTLs or 1.7%∼13.3% more eGenes at the peak (**Supp Figure 5**). It was also clear that the number of detected eGenes started to drop much earlier when incorporating highly correlated PEER factors (**Supp Figure 5**). Performing these sensitivity analyses in new studies is time-consuming and computationally expensive, especially for large cohorts with many cell types. Our results show that using the top 2000 HVGs to generate PEER factors could achieve similar power in eGenes discovery compared to using all genes while saving significant computational resources (**Figure 2B**). Furthermore, the optimal number of fitted PEER factors is not solely dependent on sample size but on how much variation can be explained. For CD4_SOX4_ cells, the inferred PEER factors did not significantly increase the eGenes detection power in most scenarios (**Figure 2C** and **Supp Table 1**); therefore, selecting the number of PEER factors in eQTL association just based on sample size could be erroneous.

To expand our exploration into other cell types, we tested the data from Neavin *et al*.^10^, who noted that the number of detected eGenes dropped with the incremental increase of PEER factors in the four iPSC clusters but not in the six fibroblast clusters (*Figure S20* in the original paper). Strong correlations among PFs were observed in four iPSC subtypes (after the 4^th^ or 5^th^ PEER factor) but not in fibroblast subtypes (**Supp Figure 6**). In the case of iPSC subtypes, fitting more PEER factors in the eQTL association analysis added more noise, which led to the loss of power. We hypothesise that the difference is due to the sample size since the input expression matrices were already quality controlled using quantile normalisation and z-transformation. There are rules of thumb for the minimum sample size required for factor analysis^16,17^, which suggest 3-20 samples per factor. When the sample size is too small, the first several PFs could explain almost all the variations, and there is not enough variation that the additional factors can explain. Thus, the following factors become strongly correlated due to overfitting (observed as very similar or even equivalent weights for certain PEER factors). The sample sizes were 79 for fibroblast and 31 for iPSC; thus, iPSC is more likely to suffer from sample size bias. We validated our hypothesis by down-sampling the fibroblast dataset (*N* = 31 to match with iPSC; **Methods**). The mean of pair-wise correlations among 10 PEER factors ranged from 0.11 to 0.99 in the six fibroblast subtypes (**Supp Figure 7**), indicating that insufficient sample sizes could result in high correlations among PEER factors even if the expression matrices were well normalised. We also down-sampled the fibroblast clusters to 40 and 50 separately and found a negligible correlation among inferred PEER factors when *N* = 50 but moderate correlations (0.004-0.39) when *N* = 40, suggesting that we might need at least five samples per factor in such a dataset.

Our results demonstrate that generating PEER factors requires careful consideration in single-cell data. The impact of how many PEER factors are included to improve the eGenes discovery power varies across different cell types. We recommend testing the correlation among inferred latent variables (also with the known covariates) and conducting sensitivity analysis to select the optimal number of latent variables to be incorporated in eQTL mapping for each cell type. As we are moving towards the era of identifying single-cell, context-dependent, and dynamic eQTL^18-20^, learning latent variables directly from single-cell level data^21,22^ and comparing them with those from pseudo-bulk would provide insights into the genetic control of gene expression at a more refined resolution.

## Conclusions

Applying methods designed for bulk RNA-seq data to pseudo-bulk data could be challenging as the assumptions might not be fully satisfied. This work highlights the pitfalls when learning PEER factors for pseudo-bulk data and presents diagnostic guidelines of performing further QC and normalization on the pseudo-bulk matrix to avoid strong and spurious correlations among the inferred factors. Optimisation for the number of PEER factors included in the eQTL association model should be carried out by a data-driven approach and using highly variable genes to generate PEER factors could achieve similar eGenes discovery power as to using all genes.

## Methods

Three single-cell datasets were used in this study to explore the performance of the PEER method. The OneK1K consortium^12^ is a population-scale single-cell RNA-seq dataset collected in Tasmania, Australia. This cohort includes 982 individuals, each with gene expression profiling for ∼1,000 (mean = 1297.0, standard deviation = 23.6) peripheral blood mononuclear cells (PBMCs). The data was quality controlled, normalised and variance stabilised by the *sctransform* method, and classified into 14 cell types (see more details in ref^12^). We further identified two individuals with problematic metrics during the preliminary test of PEER factors (one with a deficient number of cells and the other with abnormal cell composition) and removed them in the primary analysis. The sample sizes for 14 different cell types range from 795 to 980 (**Supp Table 1**). Neavin et al.^10^ collected 64,018 fibroblasts from 79 donors and 19,967 iPSC from 31 donors. The fibroblast data were classified into six subtypes and iPSCs into four subtypes. For each subpopulation, the pseudo-bulk was calculated as the mean expression per gene per individual and then quantile-normalised and z-transformed.

PEER factors are latent variables that can explain the variability in gene expression. The original method^3^ was proposed in 2010, and the software^4^ was released in 2012. We used the R package ‘peer’ (v1.0) to generate the PEER factors for the pseudo-bulk data applying max iterations = 2000 and the number of PEER factors = 50. Rank-Inverse Normal Transformation (RINT) was applied to the data by the function *RankNorm()* in the R package ‘*RNOmni*’^23^. The transformed matrix was standardised to a mean of zero with a unit standard deviation per gene. For analysis using the top 2000 HVGs, a refined gene list (pre-excluded genes with π_0_ > 0.9 or mean < 0.001) was ranked by their Fano factor (variance-to-mean ratio) before transformation and scaling. Note that these HVGs are not the same HVGs usually defined in the QC step of the raw expression matrix for single-cell data. The former indicates the genes with high mean variability across individuals, while the latter shows the genes that are highly variable across cells.

The eQTL association analysis was performed by Matrix eQTL (v2.3)^24^. We fit sex, age, the first six genotype PCs, and PEER factors as the covariates. The study only included SNPs located in the *cis*-region of the gene within the 1Mb from either upstream or downstream and with minor allele frequency > 5%. A local false discovery rate (LFDR) was calculated to control the false-positive rate for each chromosome tested by the R package ‘*qvalue*’^25^. An eGene was reported when at least one significant eQTL was found at LFDR < 0.05.

To investigate whether the strong correlation of PEER factors in iPSC data from Neavin *et al*.^10^ arose due to the small sample size, we randomly down-sampled the six fibroblast subtypes from 79 to 31 individuals (to match the sample size of the iPSC subtypes) 30 times and then generated PEER factors with these sub-samples. For each sub-sample, pair-wise Pearson’s correlations among 10 PEER factors were estimated. A similar down-sampling analysis was also conducted for sample sizes equal to 40 and 50.

## Code availability

The analysis code is available on https://github.com/powellgenomicslab/PEER_factors

## Acknowledgements

We thank Dr Walter Muskovic for his assistance with identifying the outliers in the OneK1K cohort. J.E.P. is supported by a National Health and Medical Research Council Investigator Fellowship (1175781). This work was also supported by National Health and Medical Research Council Project Grant (1143163), and Australian Research Council Discovery Project (190100825).

