## Supporting material for "Pitfalls and opportunities for applying PEER factors in single-cell eQTL analyses"

**Supplementary Figures**

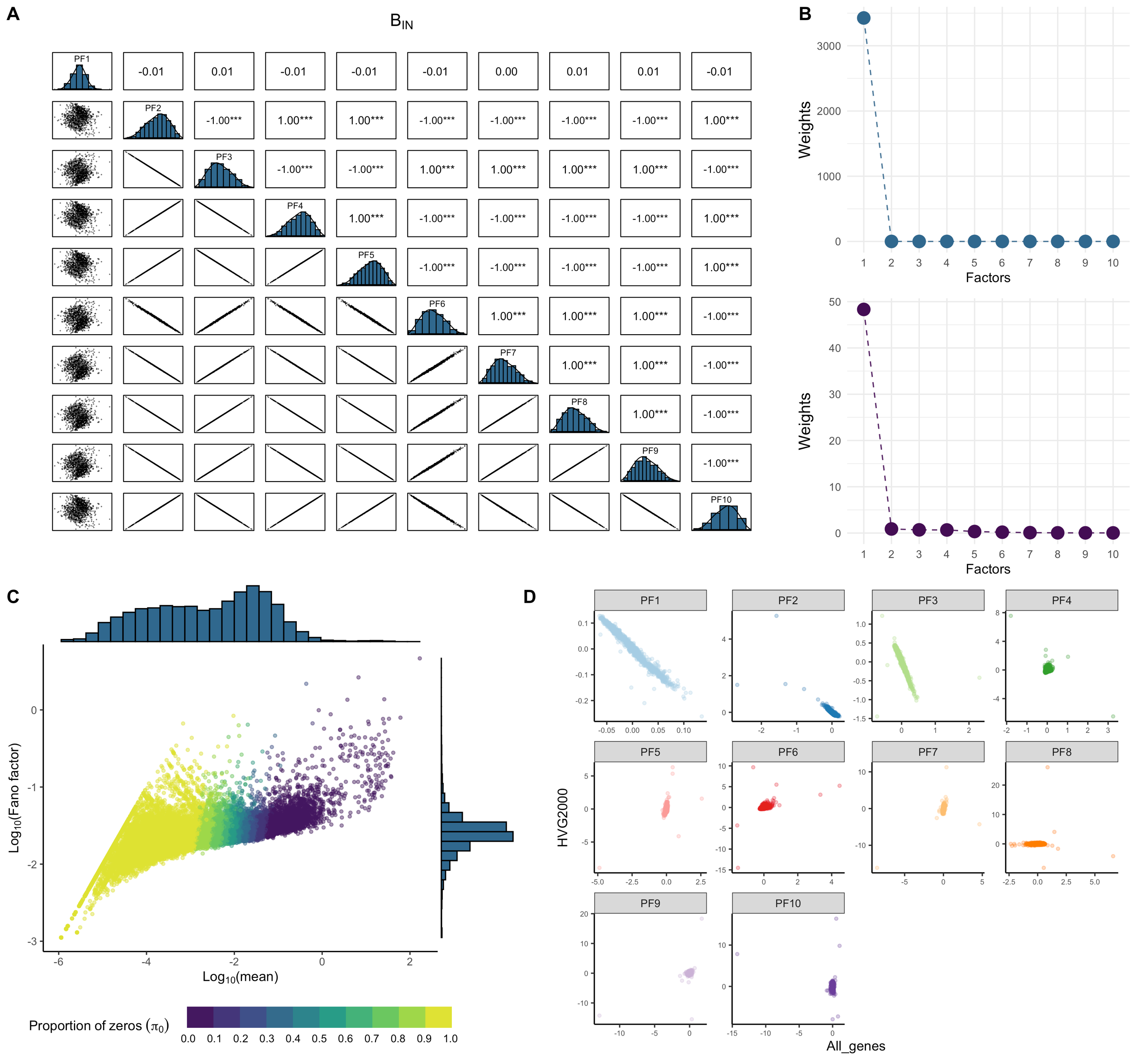

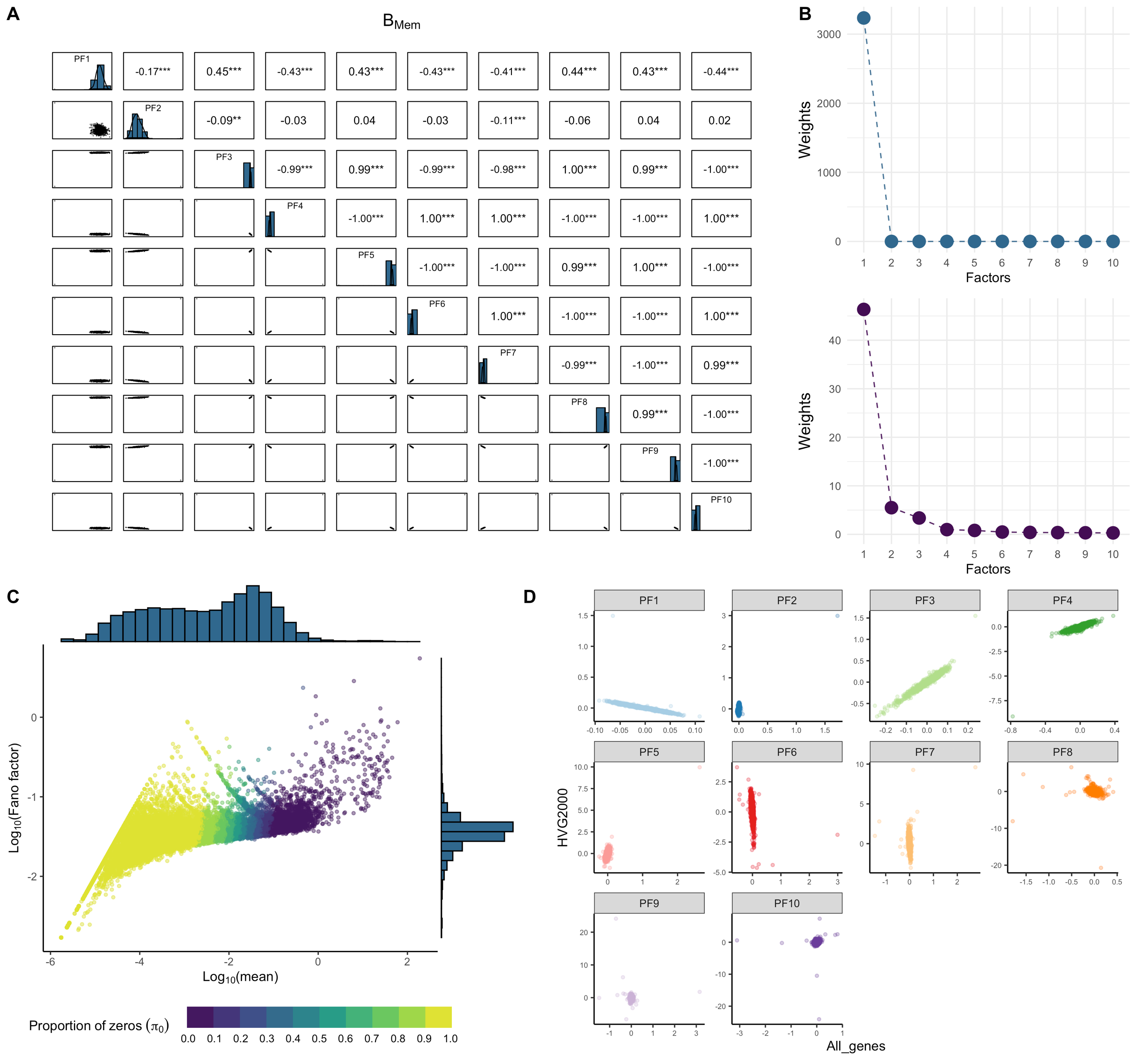

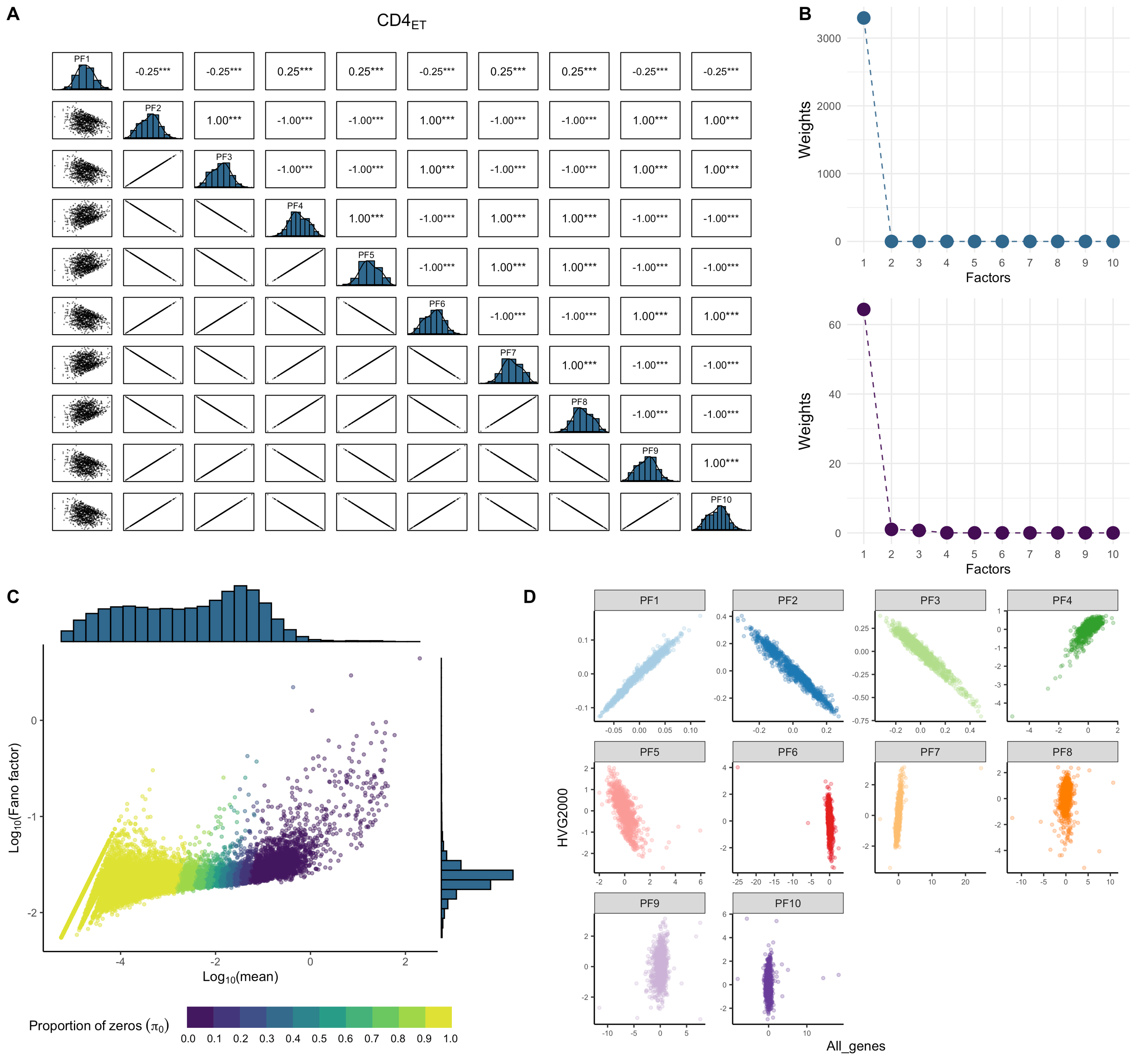

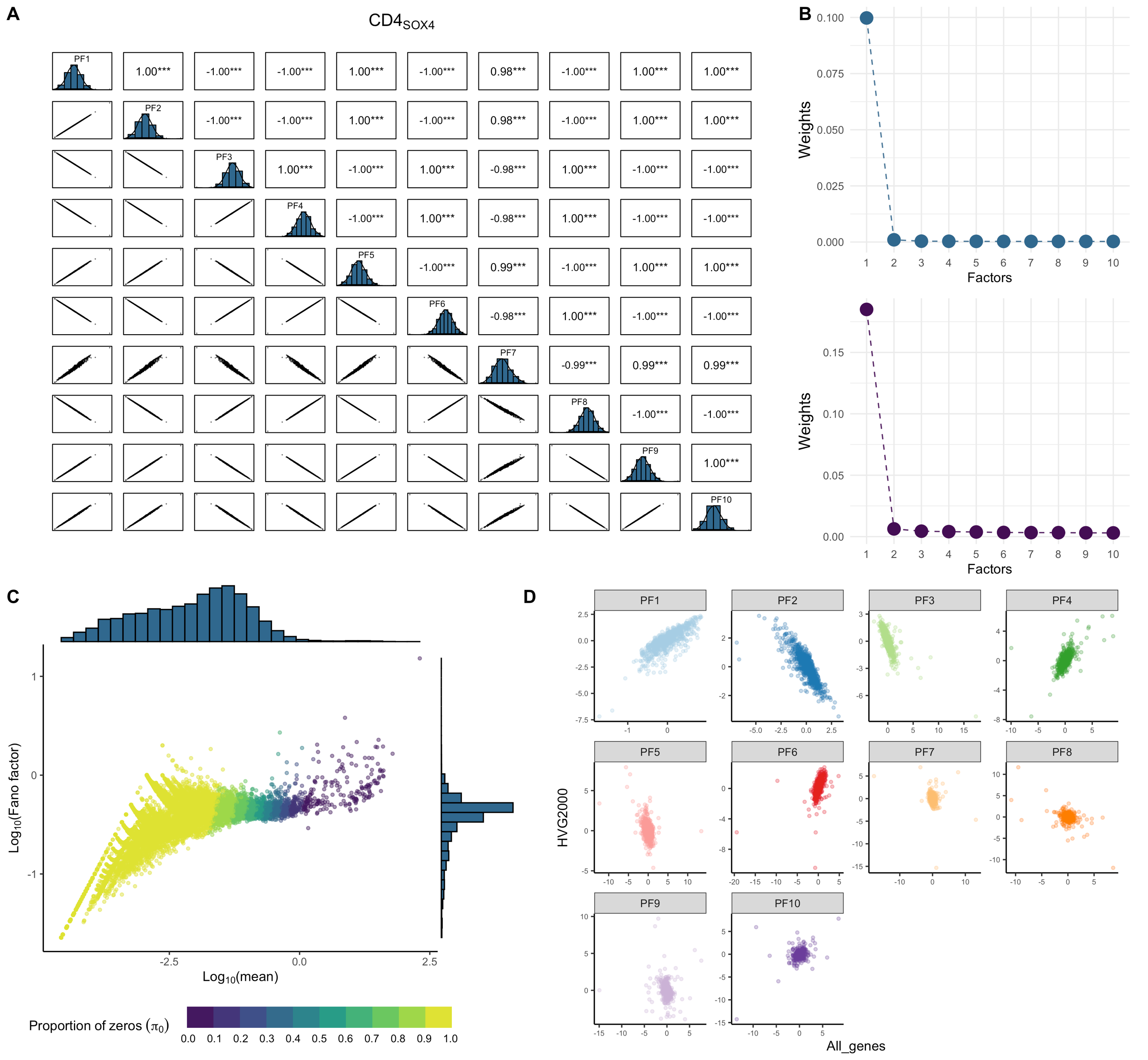

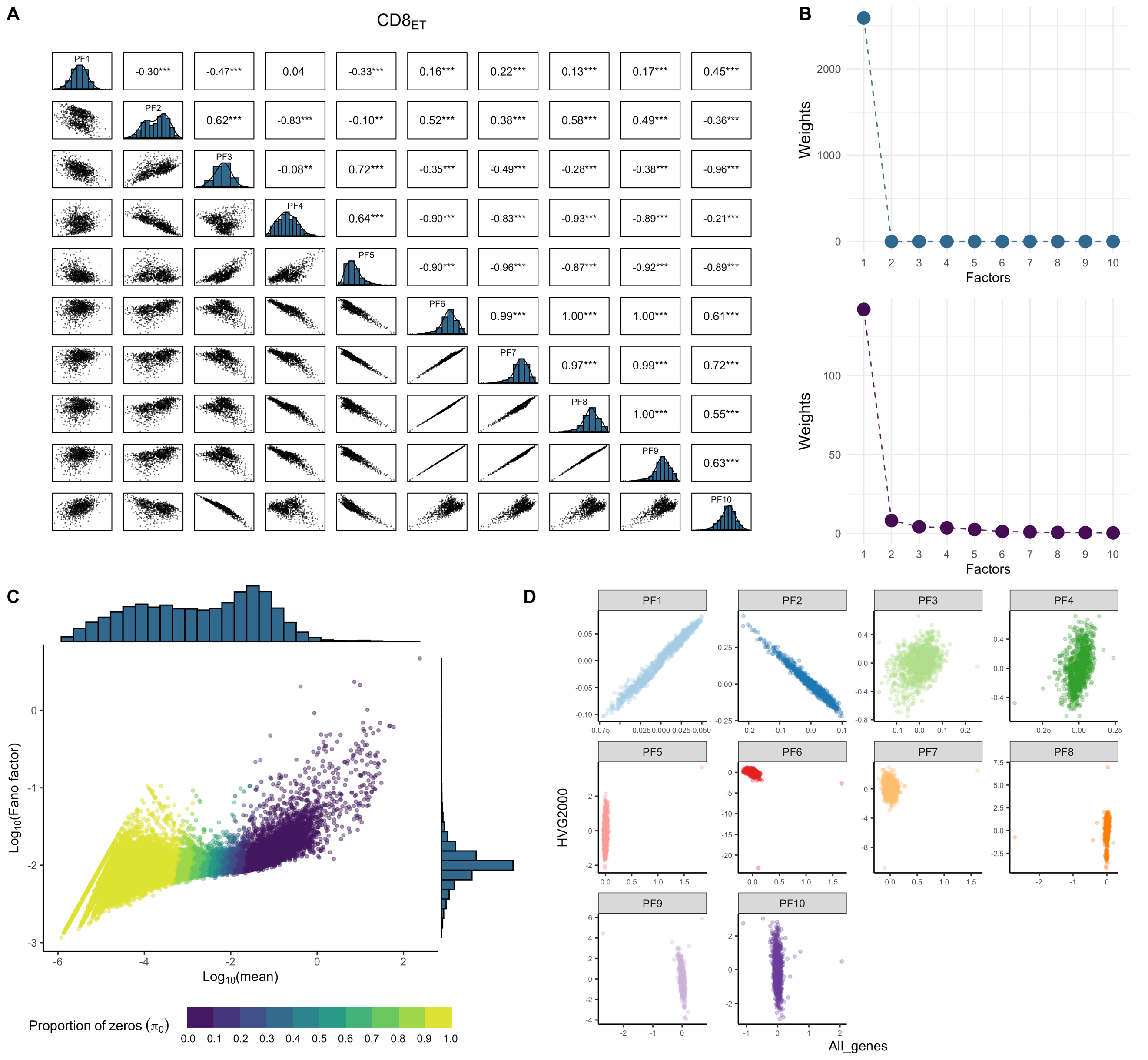

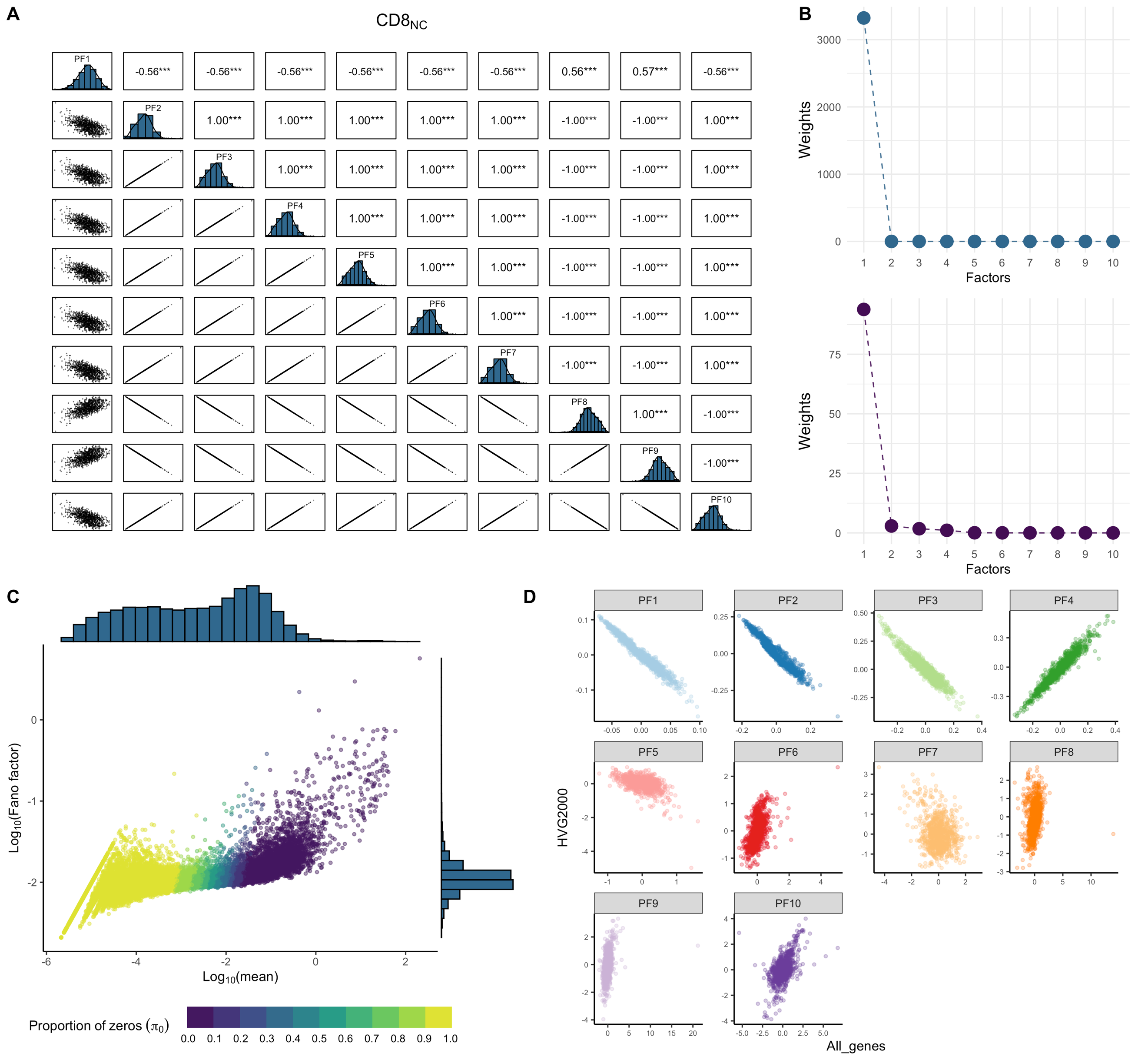

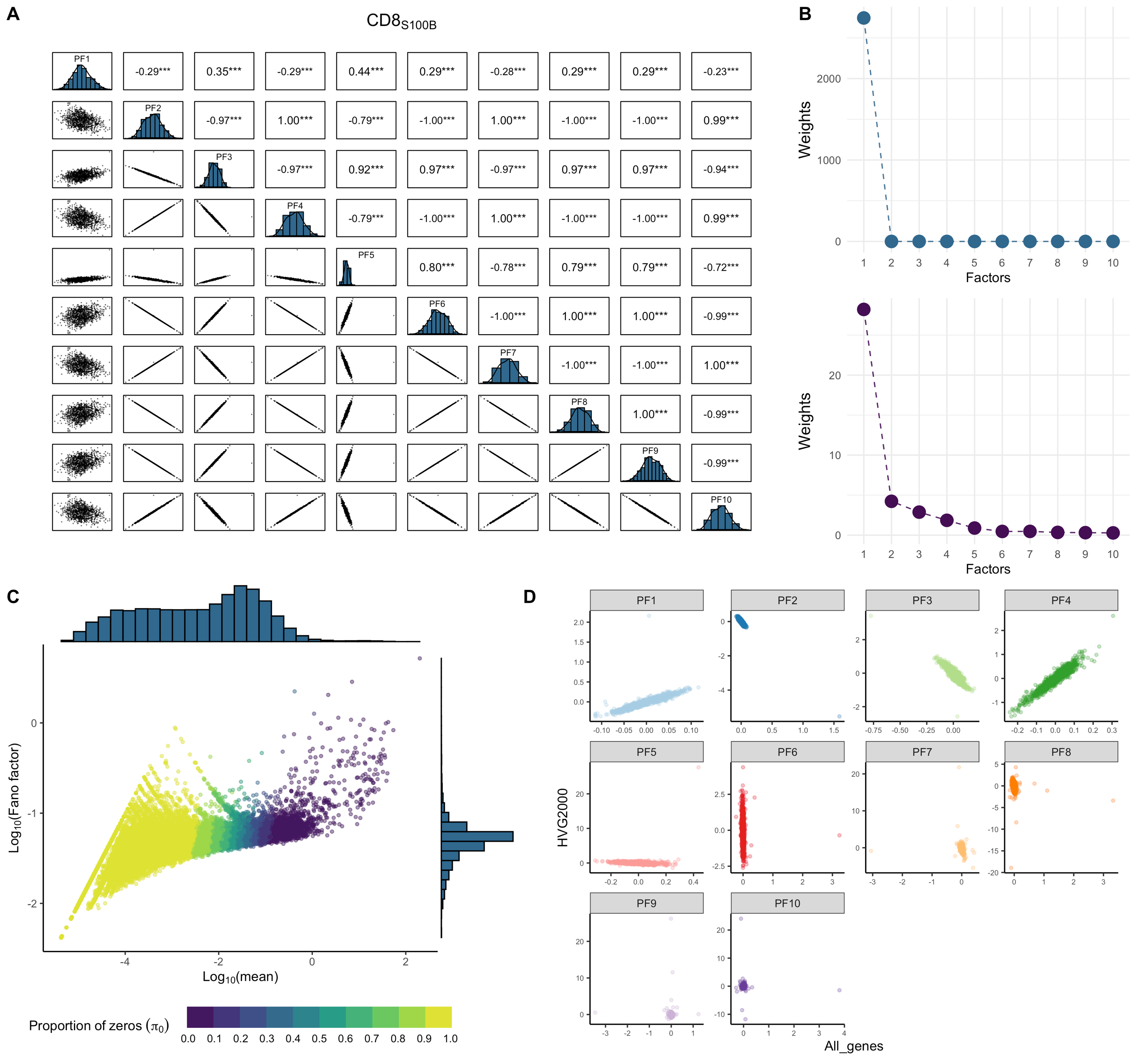

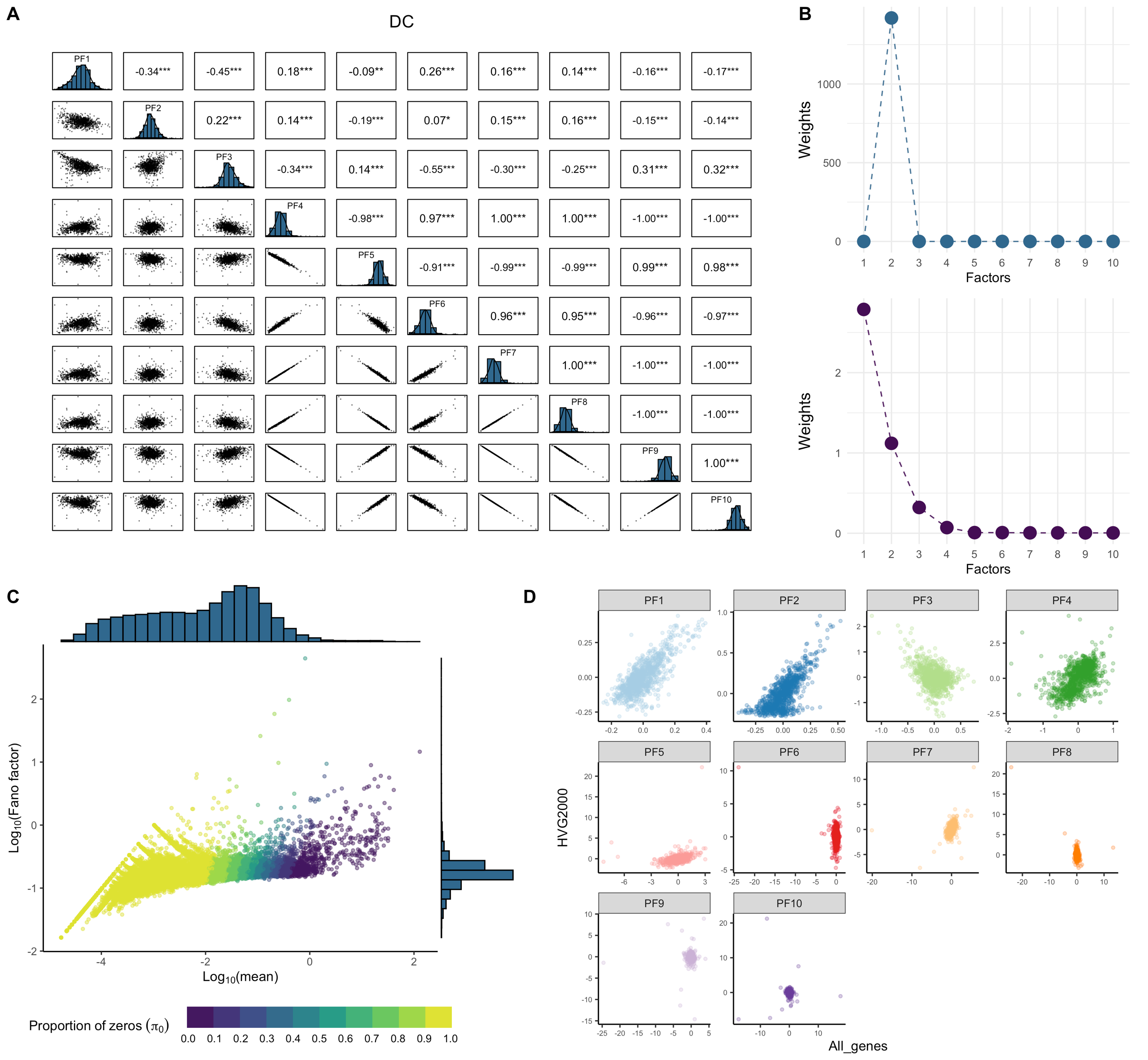

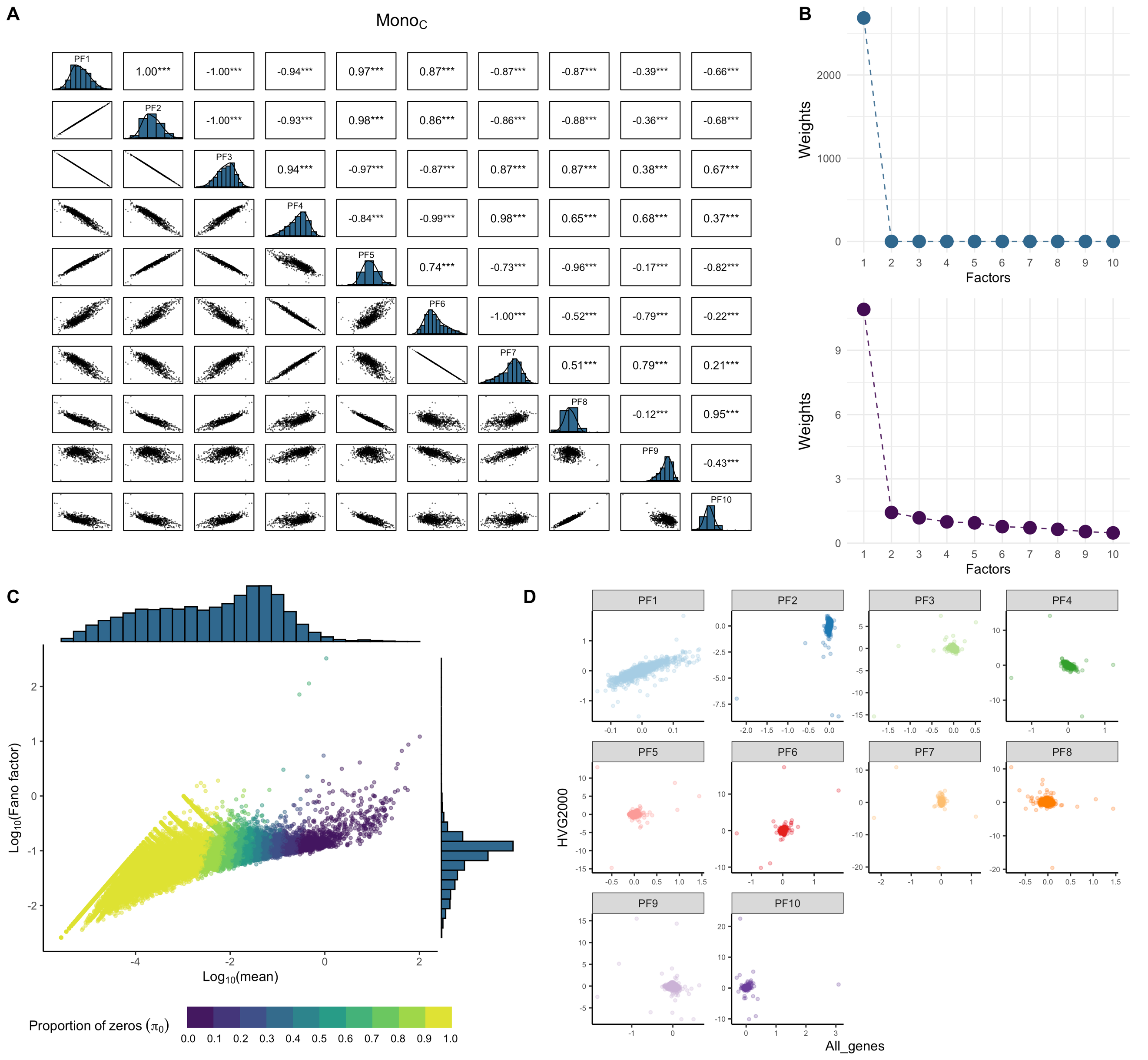

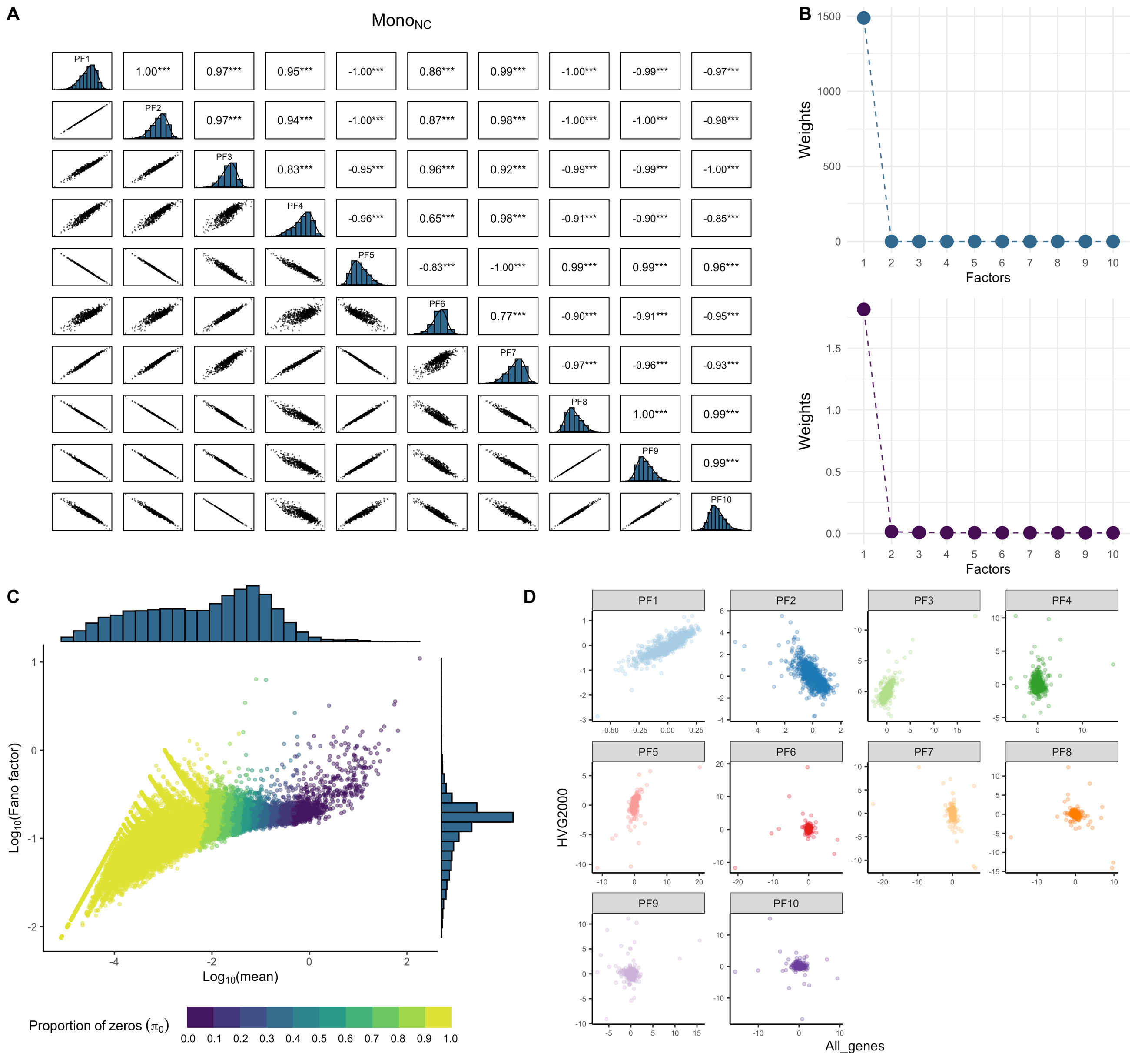

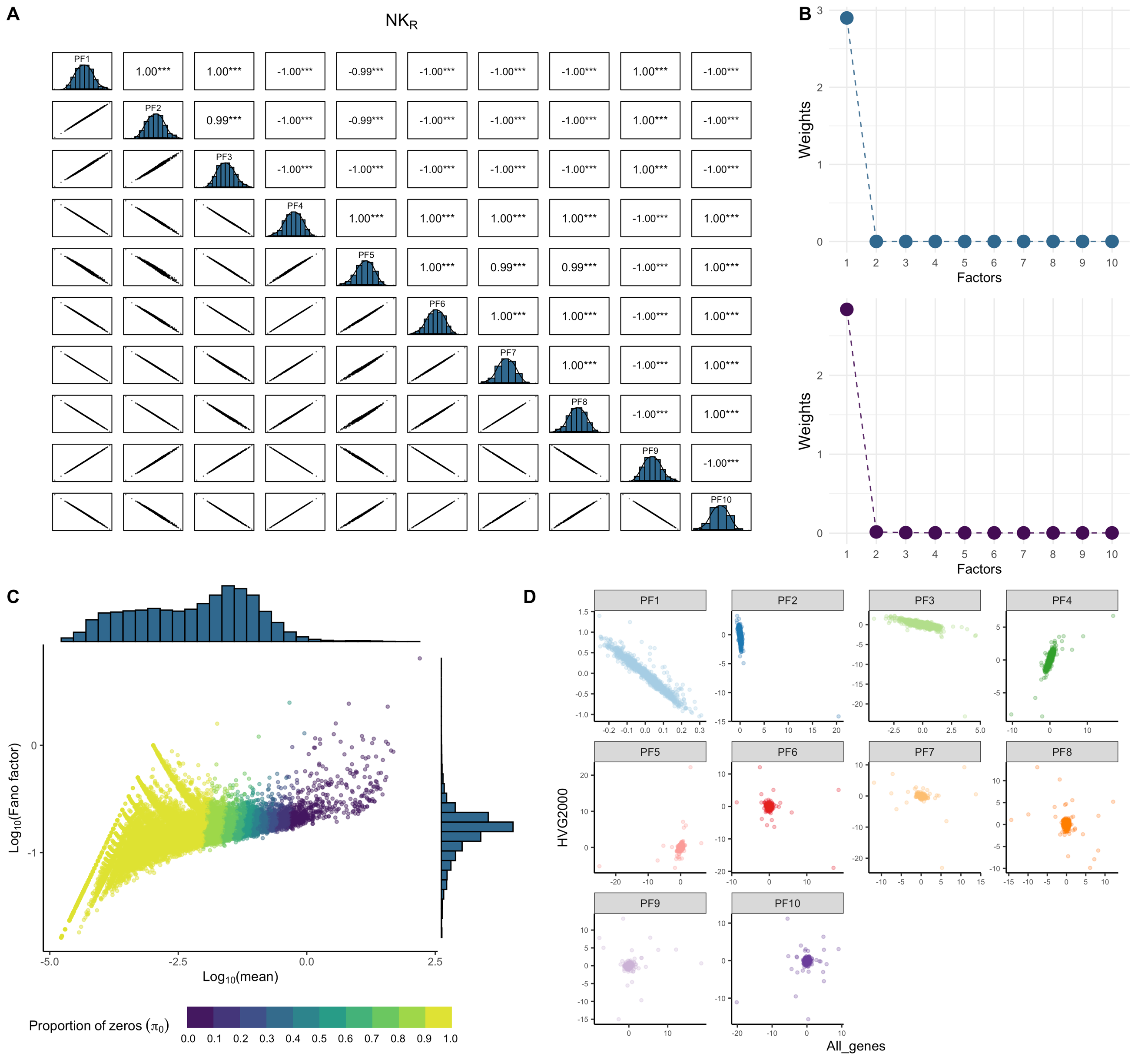

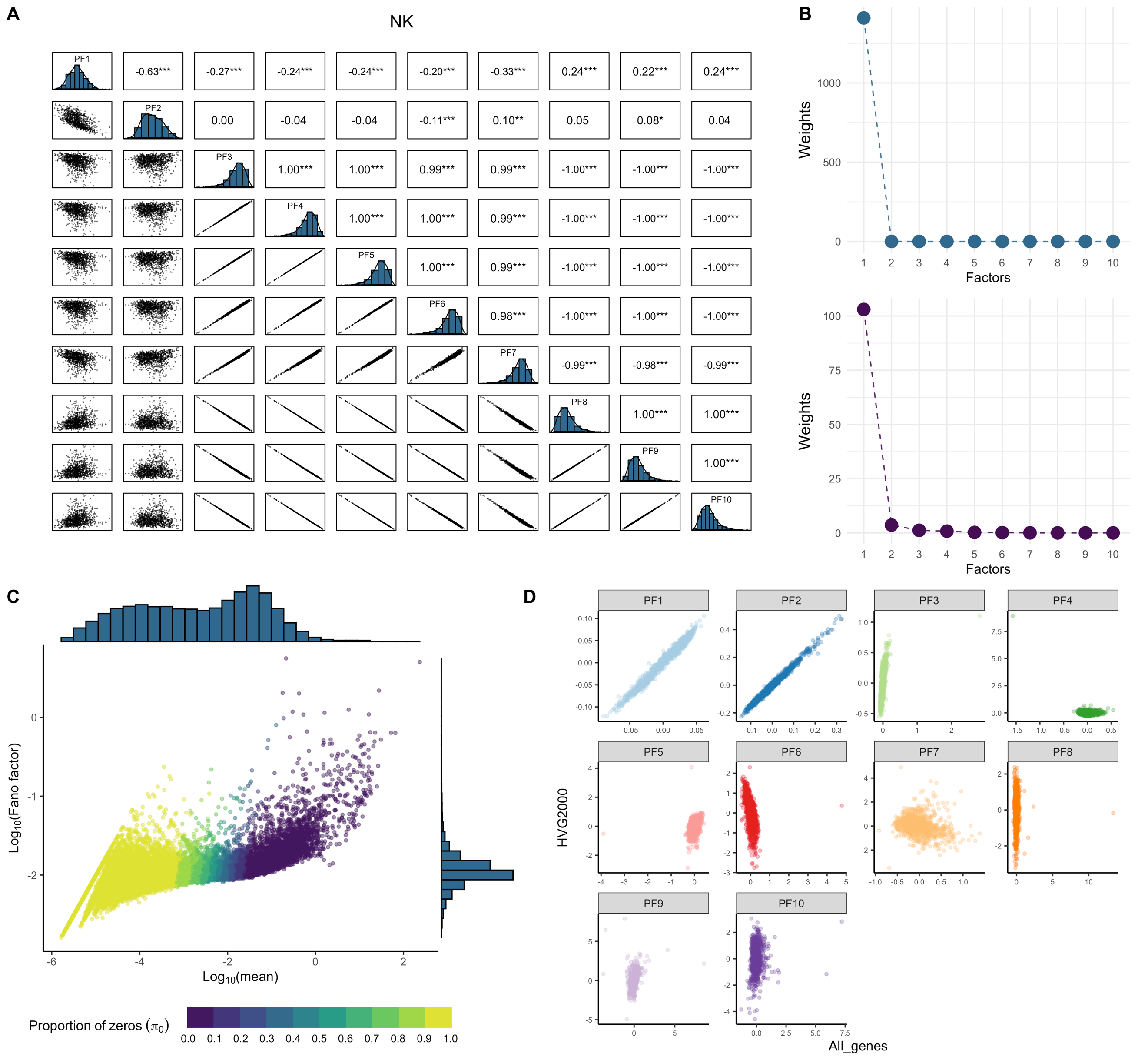

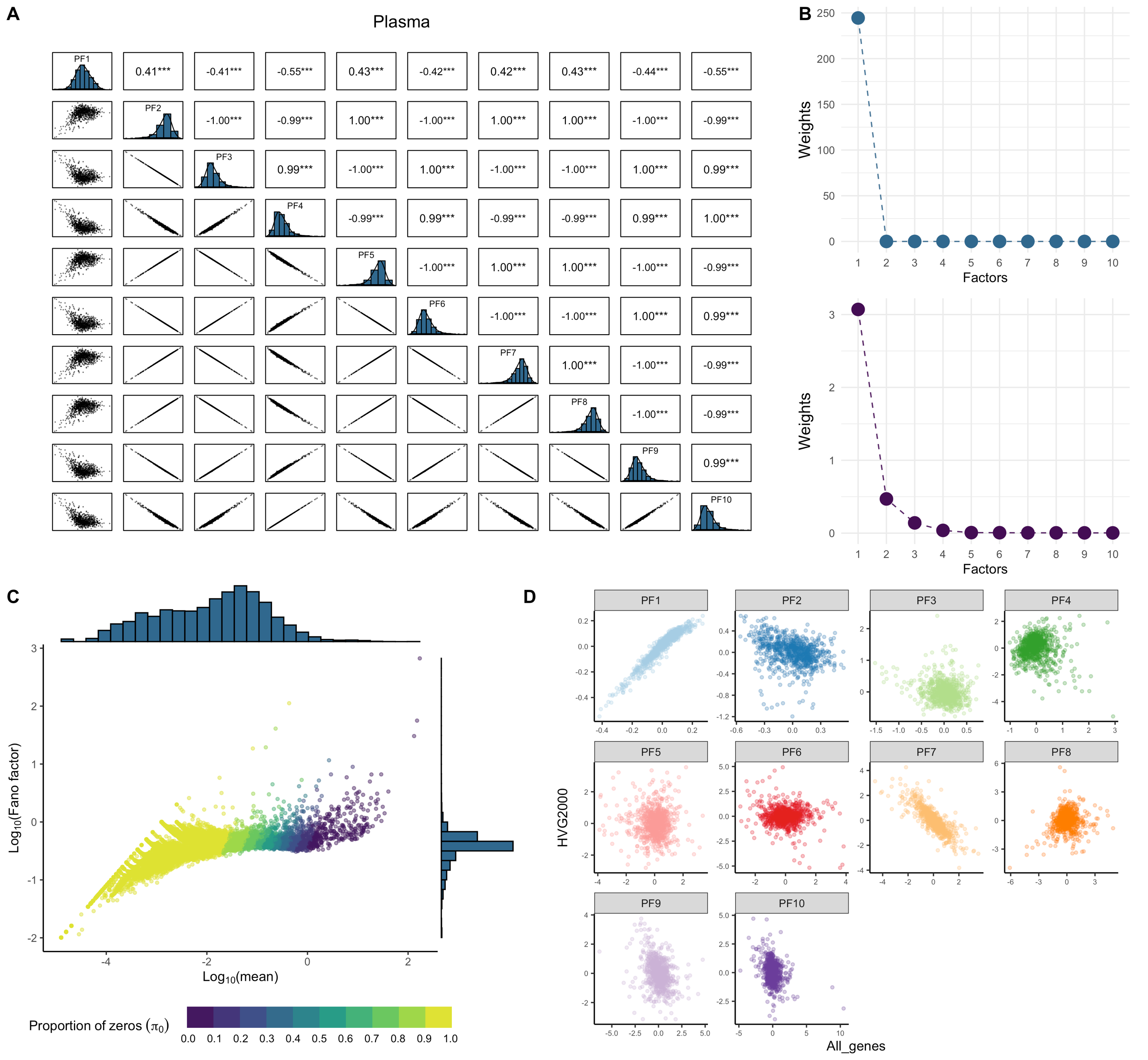

**Figure S1. Correlation among inferred PEER factors and global intra-individual mean-variance dependence**. The figure legend is the same as **Figure 1**. **A,** Pair-wise correlation plot among the first 10 PFs generated from pseudobulk expression in CD4_NC_ cells without any QC (option #1). The upper triangle panel shows the pair-wise estimates of Pearson's correlation, and the bottom triangle panel shows the pair-wise scatter plot between the PFs. The diagonal panel shows the distribution of each PF. Significance of correlation test is annotated by * *p*-value ≤ 0.05, ** ≤ 0.01, *** ≤ 0.001. **B**, Diagnostic plot of the factor weights without any further QC on the pseudo-bulk matrix (option #1, upper panel) and option #11 QC (lower panel). **C**, Relationship between intra-individual pseudo-bulk mean and Fano factor per gene. Both axes are Log10 transformed. The colour of the dots indicates the proportion of zero expression across individuals ($\pi_{0}$) for each gene. **D**, scatter plot of first 10 PEER factors generated from all genes against those from top 2000 HVGs (option #11 vs option #12). PF: PEER factor, QC: quality control, HVGs: highly variable genes.

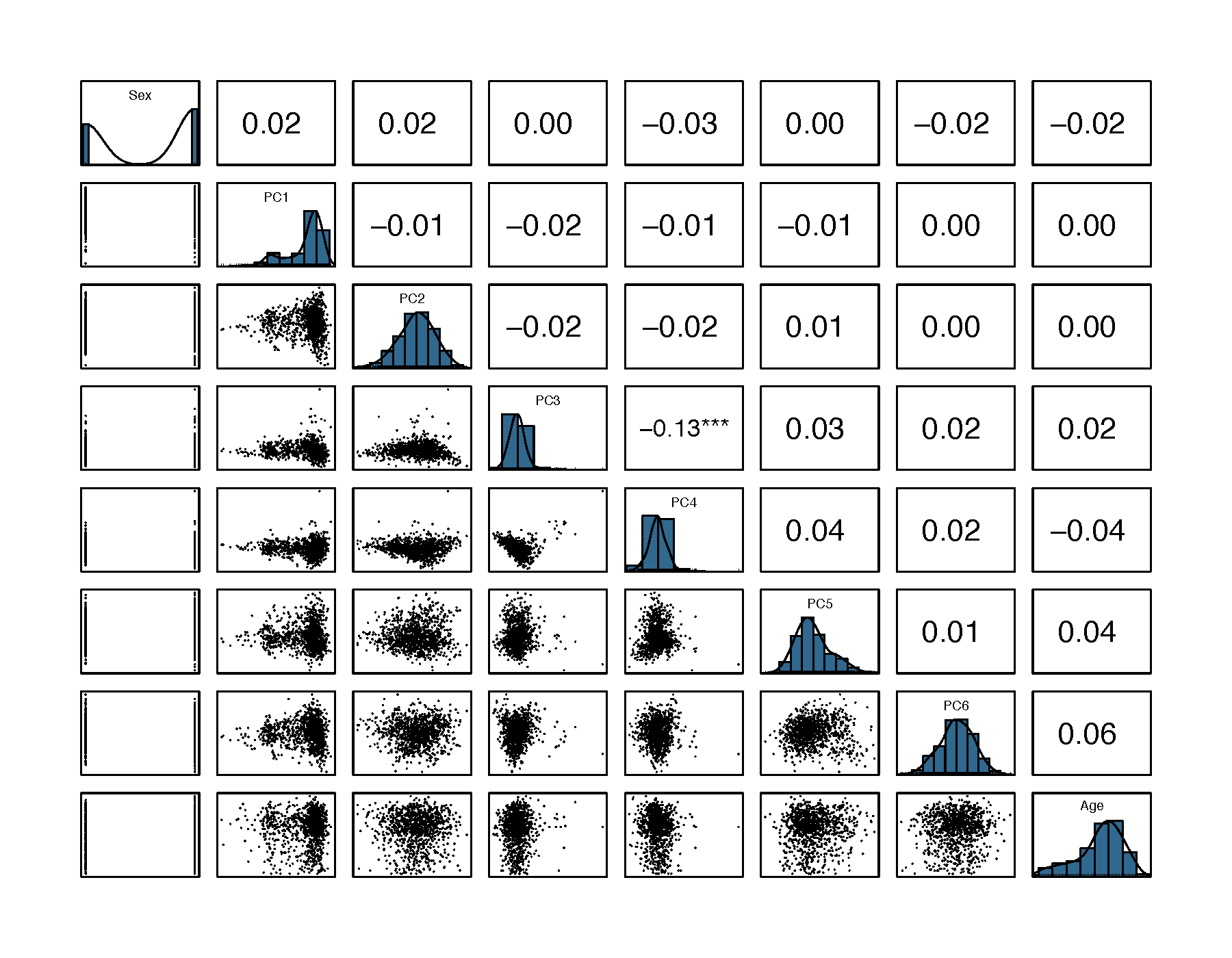

**Figure S2. Correlation among known covariates including sex, age, and first six genotype PCs**. The lower triangle denotes the scatter plot of pair-wise variables, and the red curve denotes the correlation ellipse. The upper triangle indicates the pair-wise estimate of Pearson's correlation coefficient *r*. Significance of correlation test is annotated by * *p*-value ≤ 0.05, ** ≤ 0.01, *** ≤ 0.001. The diagonal panel represents the distribution of each know covariate. PC: principal components.

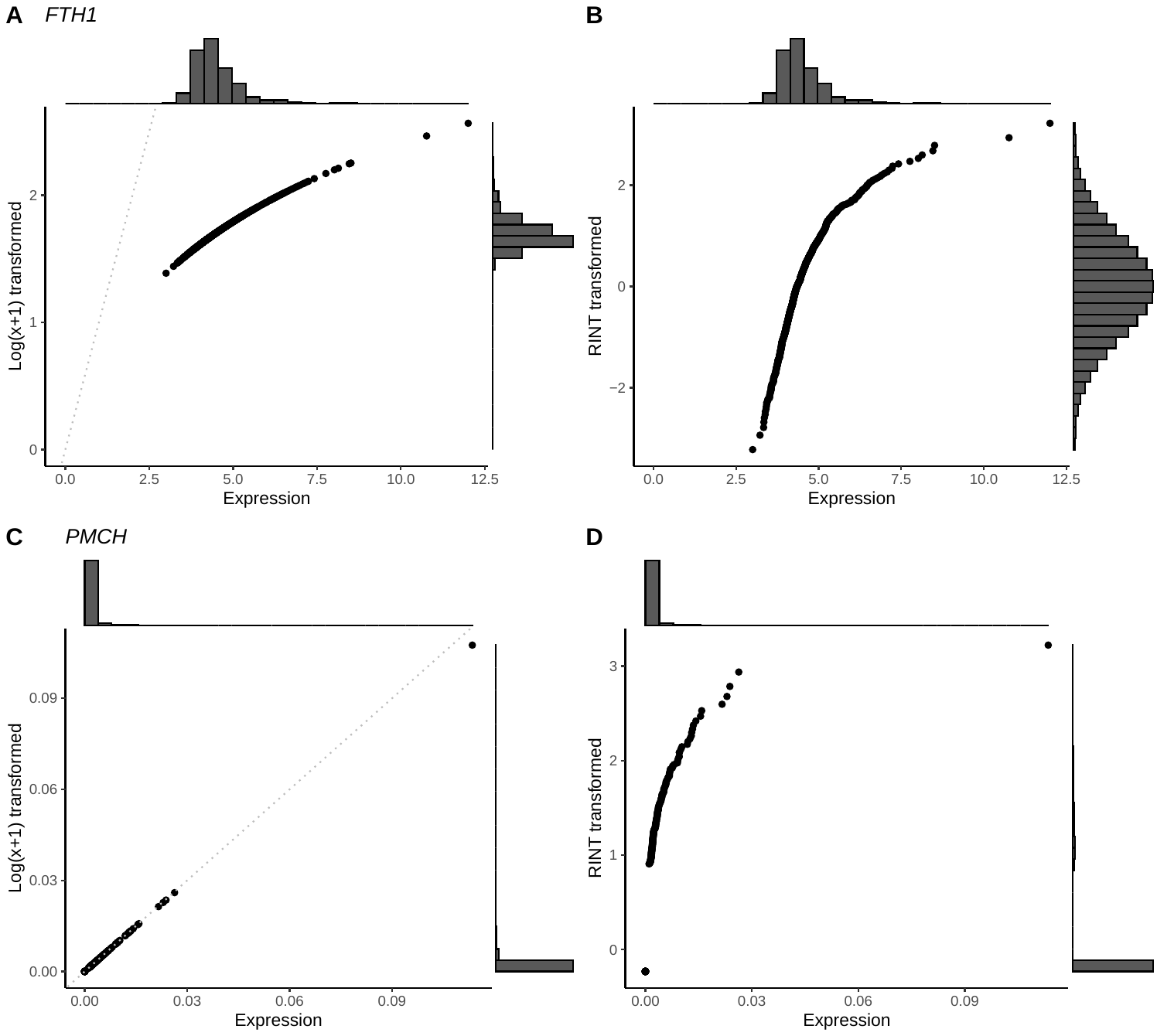

**Figure S3. Different transformation of highly expressed and lowly expressed genes**. **A-B**, x-axis is the pseudobulk expression of highly expressed *FTH1* across individuals in CD4_NC cells. The y-axis is the log(x+1) transformed or RINT transformed expression. **C-D**, The same for lowly expressed *PMCH*. The grey dashed line represents the diagonal line of the coordinate panel.

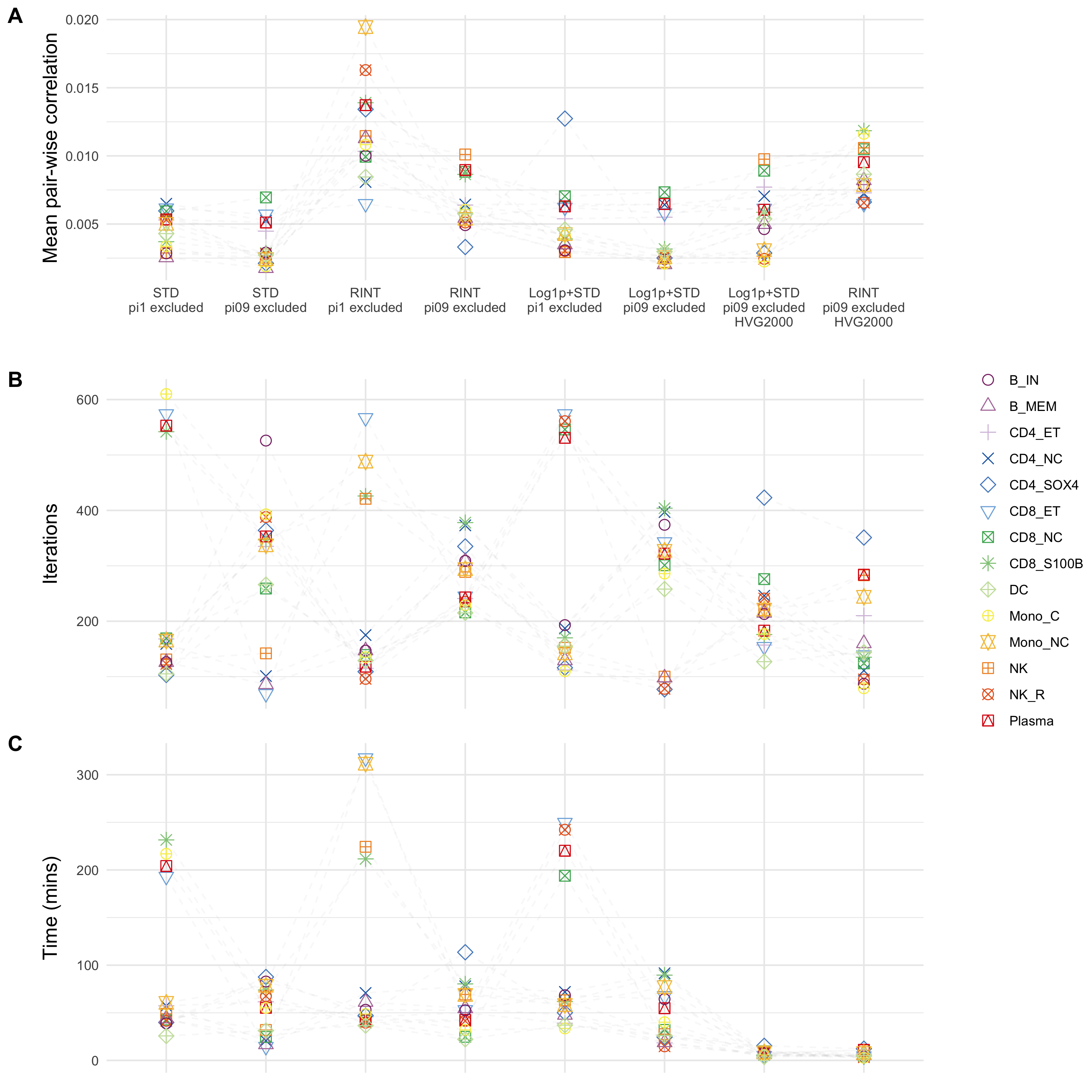

**Figure S4. Performance of the 8 candidate QC options of input matrix for PEER factor generation**. **A**, The mean pair-wise correlation among first 10 PFs. Each color and shape represent a specific cell type. The x-axis corresponds to different QC options on the pseudo-bulk matrix. **B**, Number of iterations required for the algorithm to converge. **C**, Time to generate 50 PFs.

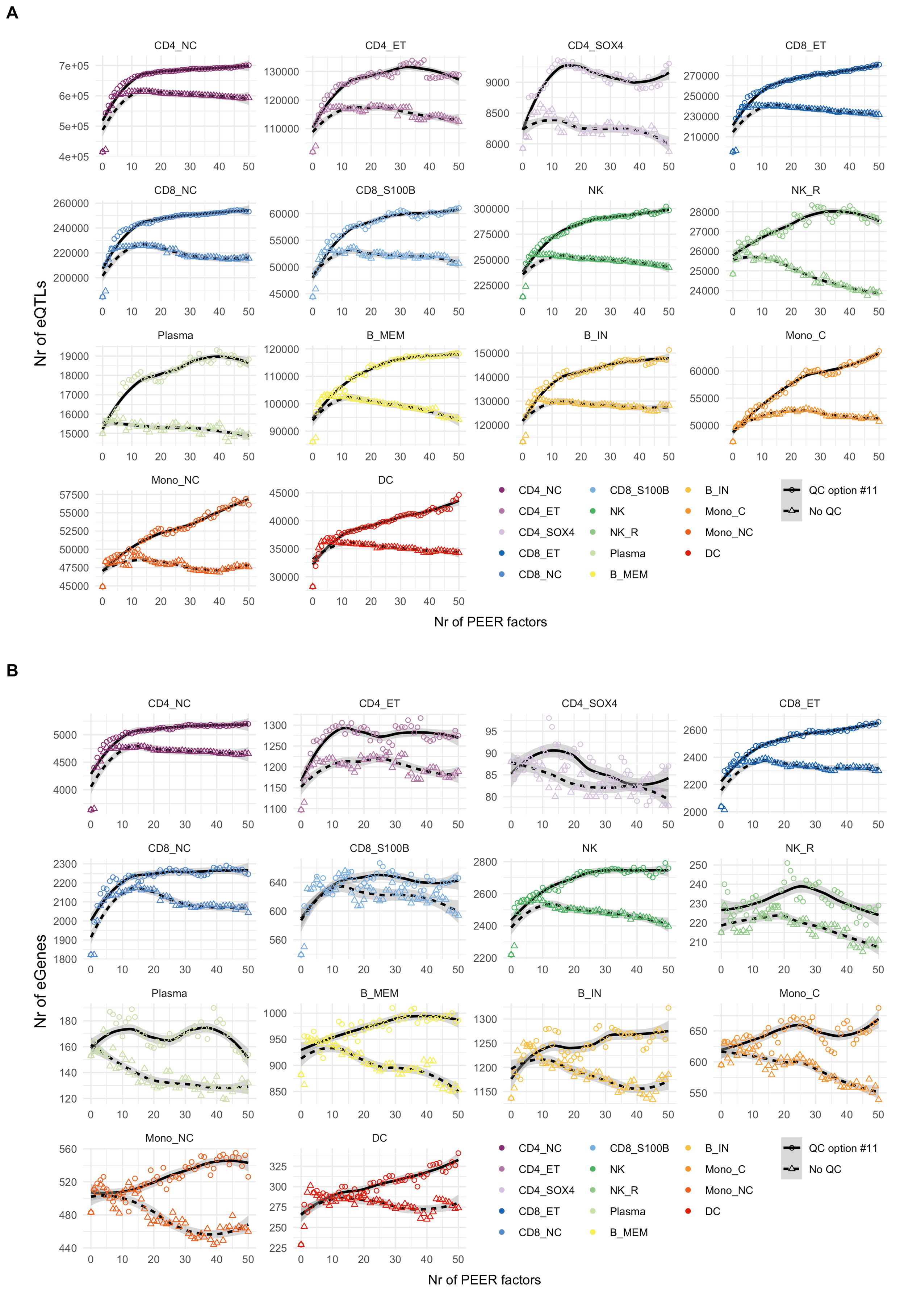

**Figure S5. Sensitivity test for eQTLs and eGenes discovery power between no QC and QC option #11**. . The y-axis represents the number of eQTLs (Panel **A**) or the number of eGenes (Panel **B**). Both x-axes denote the number of PEER factors fitted as covariates in the association model. Local regression is fitted for each QC option. The shape of each scatter point and the type of the fitted lines indicate whether using QC option #11 (circle, solid line) or no QC (triangle, dashed line) on the pseudo-bulk matrix to generate PEER factors. QC options #11 includes the steps to exclude genes with $\pi_{0}\geq0.9$, log(x+1) transformed and standardised the pseudo-bulk mean counts per gene.

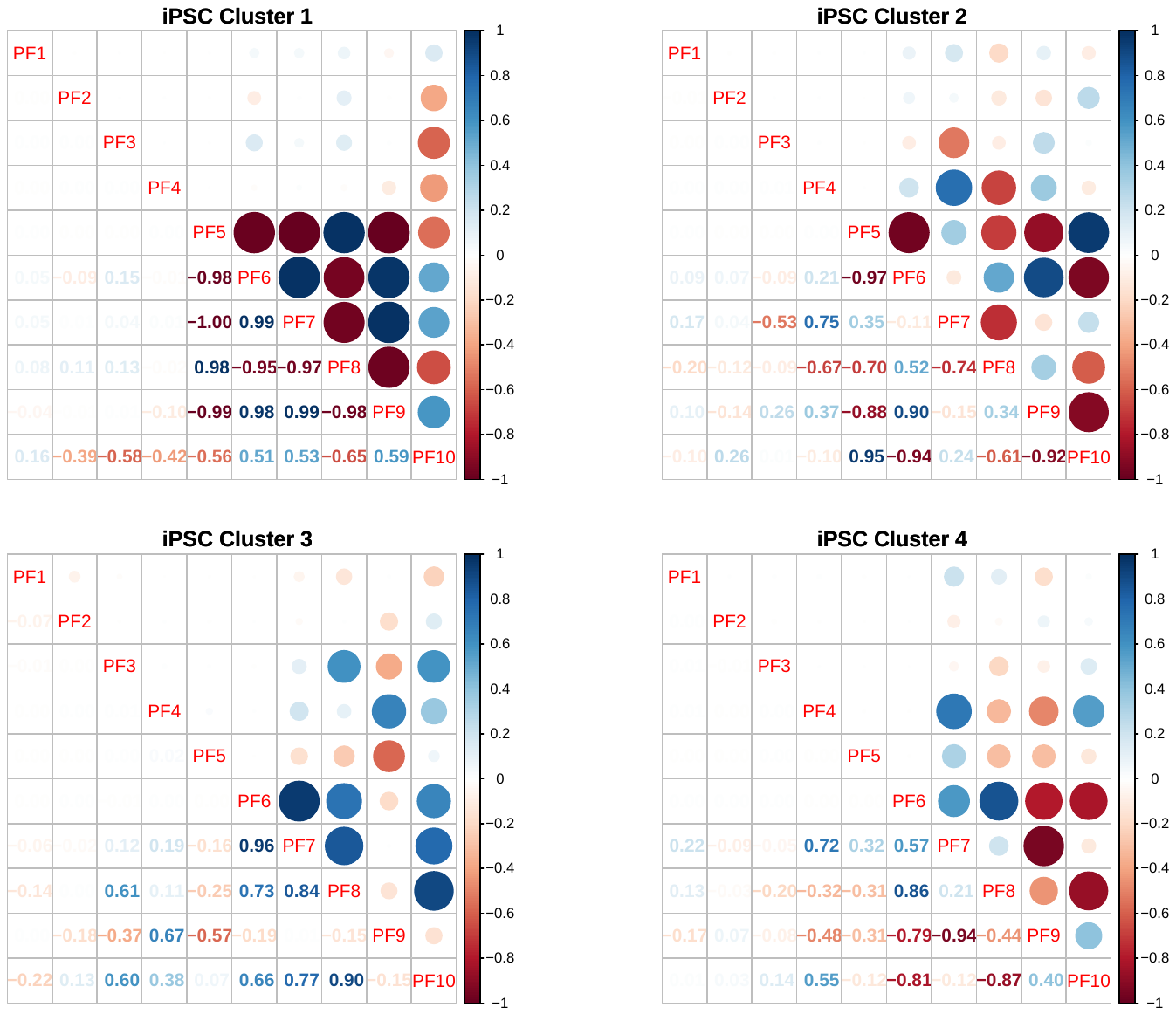

**Figure S6. The correlation plot of fibroblast and iPSC clusters from Drew et al**. Here shows the correlation plot of four iPSC clusters. The four clusters correspond to the cell population classified by the relative activity of the following regulating transcription factors (cluster 1-4 = BRF2^+^, ATF2^+^, HIC2^+^ and CEBPG^+^). The lower triangle indicates the estimate of the correlation coefficient. The circles in the upper triangle show the size and sign of the correlation by circles. The blue circle indicates positive correlation, and the red circle indicates negative correlation. The correlation in fibroblast data was negligible (most correlation coefficients < 0.003), thus was not shown here.

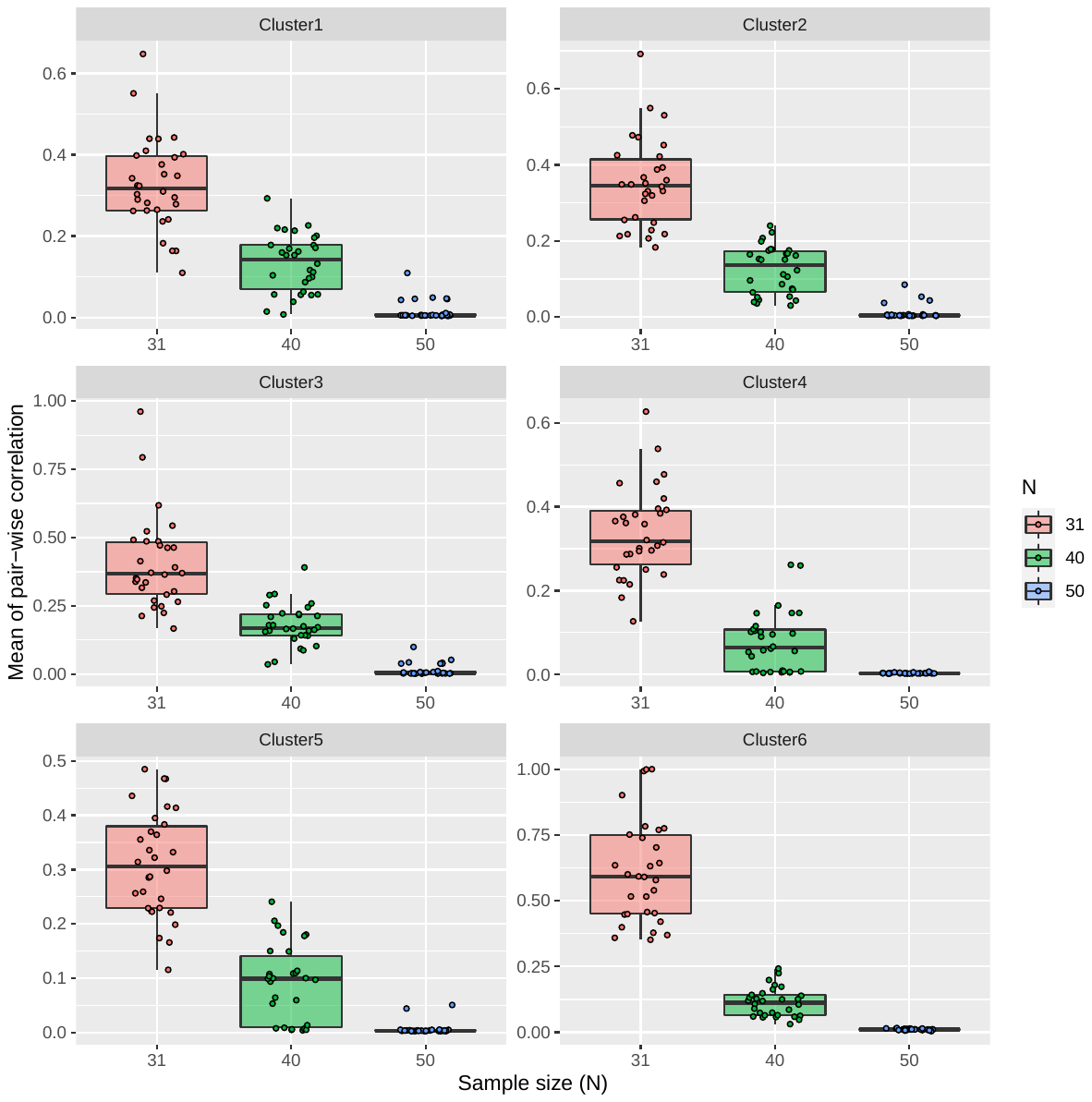

**Figure S7. Distribution of mean pair-wise correlation coefficient among PEER factors in down-sampling of fibroblast clusters**. The x-axis indicates the sample size of three down-sampling tests annotated by different color. The y-axis indicates the mean of 45 pair-wise correlations (absolute value) among first 10 PEER factors. Each dot represents one of the 30 replicates. Each panel shows one of the six clusters in the fibroblast data. The six clusters correspond to the cell population classified by the relative activity of the following regulating transcription factors (cluster 1-6 = SIX5^+^, HOXC6^+^, ATF1^+^, TEAD2^+^, KLF10^+^ and RXRB^+^).

| **Cell type** | ***N*** | **All genes**  (option #11) | | **HVG2000**  (option #12) | |
| --- | --- | --- | --- | --- | --- |
|  |  | **Prop.** | **Nr. of PF at peak** | **Prop.** | **Nr. of PF at peak** |
| B_IN | 980 | 0.165 | 50 | 0.133 | 10 |
| B_MEM | 980 | 0.145 | 29 | 0.159 | 36 |
| CD4_NC | 980 | 0.434 | 49 | 0.414 | 32 |
| CD4_ET | 980 | 0.201 | 38 | 0.209 | 28 |
| CD4_SOX4 | 857 | 0.114 | 12 | 0.034 | 20 |
| CD8_NC | 980 | 0.257 | 43 | 0.281 | 30 |
| CD8_ET | 980 | 0.304 | 50 | 0.295 | 41 |
| CD8_S100B | 979 | 0.238 | 25 | 0.230 | 30 |
| DC | 967 | 0.489 | 50 | 0.445 | 49 |
| Mono_C | 967 | 0.155 | 50 | 0.116 | 32 |
| Mono_NC | 932 | 0.149 | 46 | 0.147 | 41 |
| NK_R | 968 | 0.167 | 21 | 0.154 | 26 |
| NK | 980 | 0.258 | 49 | 0.243 | 35 |
| Plasma | 795 | 0.242 | 39 | 0.281 | 15 |

**Supplementary Table 1. The eGene detection power gain by incorporating PEER factors**. The column "Prop." indicates the power gain, and "Nr. of PF at peak" is the number of PFs fitted at the maximum number of eGenes. The power gain is measured by the proportion of new eGenes can be maximised by fitting PEER factors compared to the number when no PEER factors fitted. *N* indicates the sample size for each cell type.
